# Double-stranded RNA bending by AU-tract sequences

**DOI:** 10.1101/2020.02.17.952044

**Authors:** Alberto Marin-Gonzalez, Clara Aicart-Ramos, Mikel Marin-Baquero, Alejandro Martín-González, Maarit Suomalainen, Abhilash Kannan, J. G. Vilhena, Urs F. Greber, Fernando Moreno-Herrero, Rubén Pérez

## Abstract

Sequence-dependent structural deformations of the DNA double helix (dsDNA) have been extensively studied, where adenine tracts (A-tracts) provide a striking example for global bending in the molecule. In contrast to dsDNA, much less is known about how the nucleotide sequence affects bending deformations of double-stranded RNA (dsRNA). Using all-atom microsecond long molecular dynamics simulations we found a sequence motif consisting of alternating adenines and uracils, or AU-tracts, that bend the dsRNA helix by locally compressing the major groove. We experimentally tested this prediction using atomic force microscopy (AFM) imaging of long dsRNA molecules containing phased AU-tracts. AFM images revealed a clear intrinsic bend in these AU-tracts molecules, as quantified by a significantly lower persistence length compared to dsRNA molecules of arbitrary sequence. The bent structure of AU-tracts here described might play a role in sequence-specific recognition of dsRNAs by dsRNA-interacting proteins or impact the folding of RNA into intricate tertiary and quaternary structures.

## MAIN TEXT

Double-stranded RNA (dsRNA) plays a central role in a number of biological processes. For instance, dsRNA molecules are involved in the regulation of gene expression by RNAi (1), or in the host responses to dsRNA encoded by viruses (2) (3) (4). In addition, dsRNA helices perform key functions as an essential part of tertiary RNA structures, including tRNA and riboswitches (5) (6), and of macromolecular RNA-protein complexes such as ribosomal subunits and the spliceosome (7) (8) (9). In many of these processes, dsRNA helices are not straight, but adopt a bent or kinked conformation e.g. during the folding of RNA into complex 3D structures (10) (11) (12) or in dsRNA:protein interactions (13) (14) (15). Motivated by these findings, immense research efforts have characterized the effect of helical imperfections, such as bulges or internal loops, on dsRNA conformations (16) (17). Nevertheless, the question of whether the nucleotide sequence can induce bending in canonical, Watson-Crick base-paired dsRNA helices remains unanswered.

In contrast to dsRNA, sequence-induced bending in the canonical DNA double-helix (dsDNA) is well-characterized. A prime example of such sequence-dependent deformations are the so-called A-tracts, runs of adenines and thymines without a TA step that, when in phase with the helical pitch yield a significant global curvature of the dsDNA (18) (19) (20) (21) (22) (23) (24). Besides their bending character, A-tracts show a peculiar conformation at the molecular level, with a characteristic narrow minor groove (25). Remarkably, both the intrinsic bending induced by A-tracts and the molecular conformation of these sequences are thought to have biological relevance. The former seems to stabilize DNA tertiary structures, such as loops and supercoils (26) (27), whereas the latter is used by proteins to achieve binding specificity (28) (29).

Scattered experimental evidence suggests the existence of sequence-induced bending in a Watson-Crick base-paired RNA duplex. Early crystallographic works reported helical kinks in the structure of an RNA duplex consisting of alternating adenines and uracils (30) (31). However, this bent conformation was stabilized by the intermolecular interactions among the molecules forming the crystal and, therefore, bending could not be attributed to the RNA duplex alone. In parallel, theoretical methods, such as molecular dynamics (MD) and Monte Carlo (MC) simulations have provided valuable insight on sequence-dependent dsRNA conformations (32) (33) (34) (35) (36) (37). In fact, recent MD studies have predicted strong sequence effects on the dsRNA shape (32) (33) and flexibility (34), which could potentially lead to sequence-induced bending. Such simulation techniques hold great potential in deciphering the sequence-dependent dsRNA conformational landscape, provided that the computational predictions are thoroughly tested against experimental measurements. However, such comparison remains challenging due, in part, to the limited availability on high-resolution dsRNA experimental structures and the number of artifacts that are often found, e.g. in crystal structures (35).

Here, we present a series of steps that led us to the direct experimental observation of single dsRNA molecules bent by the nucleotide sequence. We combined MD simulations and atomic force microscopy (AFM) experiments; a technique especially suited for studying dsRNA bending, as demonstrated by AFM measurements of the dsRNA persistence length (38) (39). We first performed a systematic analysis of how the sequence affects the structure of dsRNA using MD simulations. Our simulations predicted that a sequence motif, that we named AU-tract, would cause a bend in the RNA double-helix. We then synthesized long dsRNA molecules containing AU-tracts in phase with the helical pitch. Analysis of AFM images of these molecules revealed that they were indeed significantly more bent than control dsRNA molecules of arbitrary sequence. Our work unveils the phenomenon of sequence-induced bending in dsRNA, challenging the traditional picture of dsRNA as an invariant double helix.

## Results and Discussion

### The dsRNA sequence affects the width of the major groove, the extension, and twist of dsRNA

In order to explore how the nucleotide sequence affects the dsRNA structure, we first analyzed a set of six MD simulations from a previous work (34). These simulations had been performed on benchmark dsRNA sequences of the form G_4_(NN)_8_G_4_, with NN=AA, AC, AG, AU, CG, GG (**Table I**), where the G_4_ regions in the termini had been included to prevent edge fraying and were excluded from the analysis. We measured the size of the grooves of these benchmark sequences using the software Curves+ (40) and found that the major groove width was highly dependent on the sequence, being able to change by as much as 6 Å. This is shown in **Fig. 1a**, where we represent the values of the major groove width along the helical axis for the benchmark molecules. Notice that, because these sequences consist of repeating dinucleotides, their major groove should be regular along the helical axis, as manifested by the flat lines of **Fig. 1a**. In contrast, the minor groove dimensions as well as the major groove depth did not significantly change with the sequence (**Fig. S1**).

**Table I.**
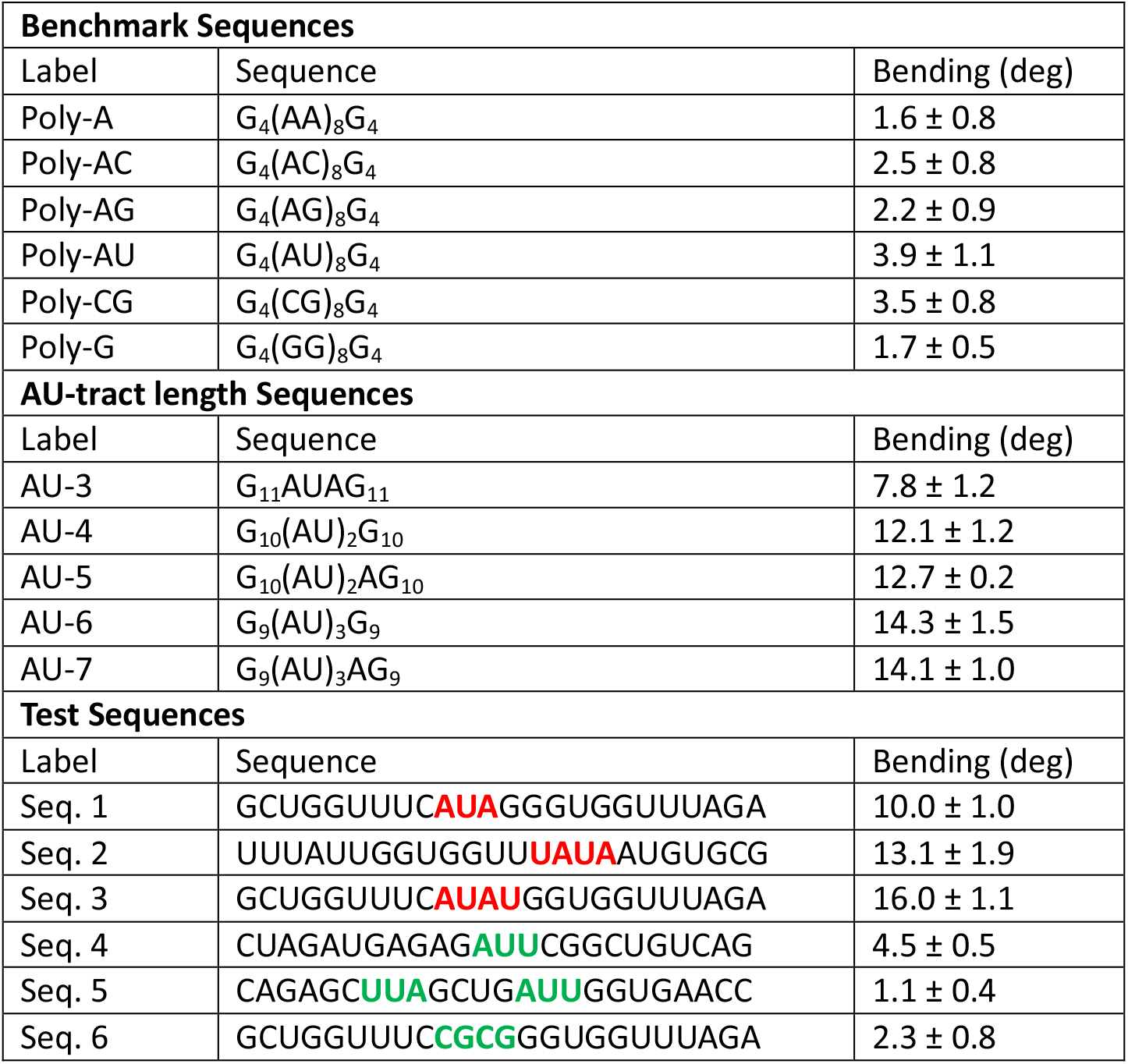
dsRNA sequences studied in this work by MD simulations. dsRNA sequences are represented in abbreviated form without the complementary strand and written from the 5’ end to the 3’ end. Benchmark sequences were selected from a previous work (34). These sequences consisted of 8 repetitions of dinucleotides flanked on both sides by G_4_. AU-tract length sequences were of approximately the same size (24 or 25 bp, depending on whether the AU-tract comprised an even or odd number of base pairs) and contained an AU-tract exactly at the center of the sequence. These centered AU-tracts were of varying lengths from three (AU-3) to seven (AU-7) base pairs. Test sequences were designed to include AU-tracts in different contexts (highlighted in red, Seqs. 1-3) and to contain other potential bending motifs, such as AUU, UAA or CGCG (highlighted in green, Seqs. 4-6). In order to calculate the bending angle, we split the trajectory into five windows of 200 ns and obtained the average structure of each window. We then computed the bending angle of these average structures using Curves+ and neglecting four base pairs in each of the termini of the molecule. The final value of the bending angle is the mean of the measurements of these windows and the error is the standard deviation.

**Figure 1.**
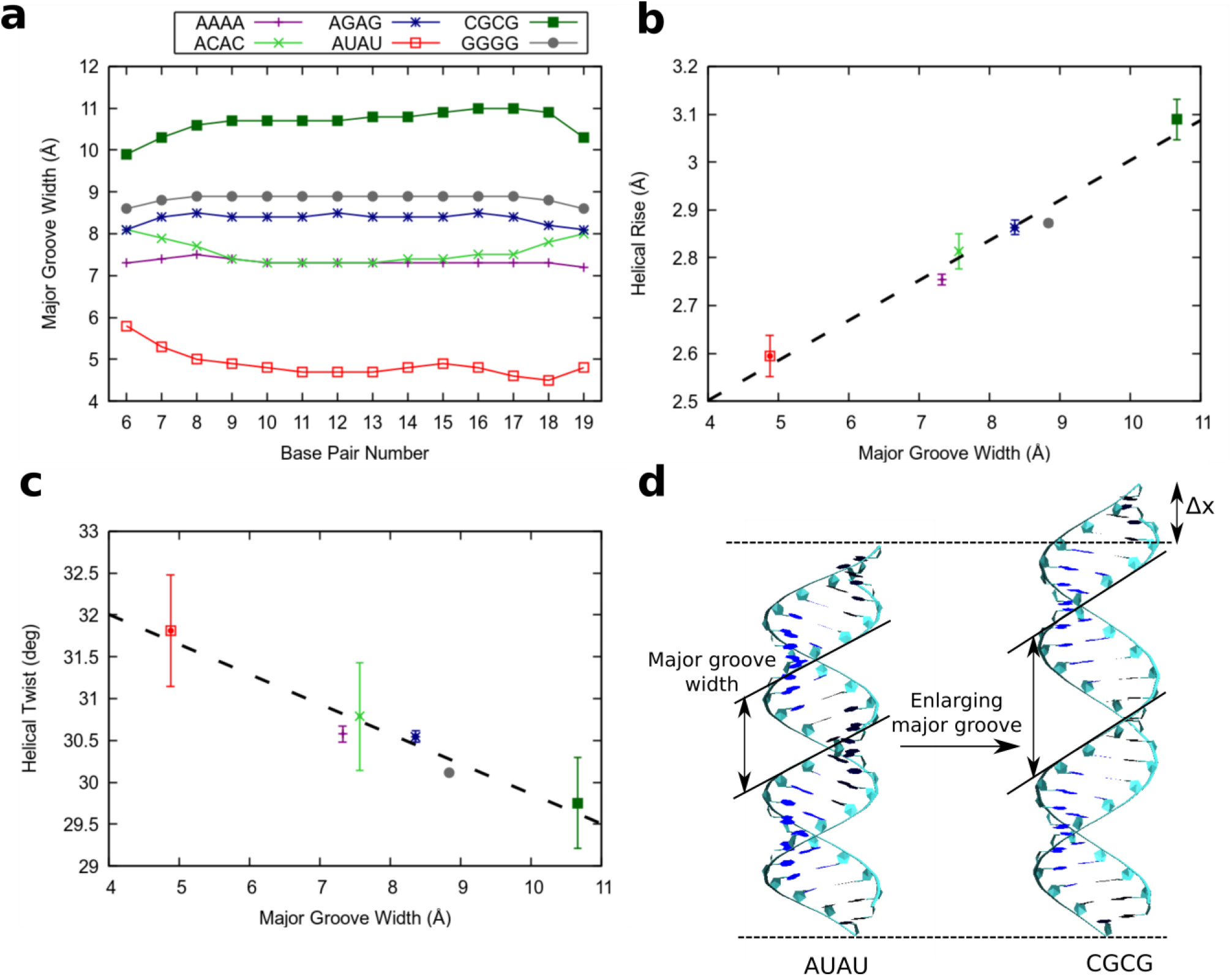
The major groove width modulates the overall structure of the dsRNA helix. **a**, Values of the major groove width measured along the helix for the benchmark sequences (Table 1). The average structures over the simulation time were computed for the benchmark molecules using the software AmberTools (63). These structures were then analyzed with the software Curves+ (40) to obtain the values of the major groove width along the sequence. **b**, Mean helical rise and **c**, helical twist of the benchmark molecules measured as a function of the major groove width. Values of the helical rise and helical twist were obtained for each base pair step from the average structures of the benchmark molecules using the software Curves+ (40). These values, together with the major groove width values from panel a, were then averaged over the 15 central base pair steps. The legend is the same as in panel a. Error bars are the standard error of the mean. X-axis error bars are within the symbols. The dotted line represents a fit of the data to a linear function. **d**, Average structures of the poly-AU and poly-CG over the simulation time. These structures illustrate how the sequence induces an elongation in the molecule by enlarging the major groove.

Notably, the major groove width was primarily responsible for modulating the extension and number of turns of the molecules. This was quantified by means of the helical rise and helical twist parameters, which were highly correlated (R=0.987) and anticorrelated (R=−0.974), respectively, with the major groove width (**Fig. 1b, c**). These results showed that the nucleotide sequence could simultaneously induce an elongation and unwinding of the dsRNA by expanding the major groove. Conversely, contraction of the major groove resulted in shrinkage and overwinding of the double-helix. This mechanism is illustrated in **Fig. 1d**, where we computed the average structures over the 1 μs simulation time of the two sequences with extreme values of major groove width: the poly-AU and the poly-CG. The former was the most compact sequence with a very narrow major groove. As the major groove was enlarged, the molecule approached a stretched and unwound conformation, which was maximal with the poly-CG sequence. The helical rise of the poly-CG and the poly-AU molecule was ± 10% of the canonical value of the extension per base pair of dsRNA, which is ~2.9 Å (41). Similar results for the sequence dependence of the major groove width and the high correlation (anticorrelation) with the helical rise (twist) were obtained when analyzing our data with the alternative 3DNA software (42) (**Figs. S2, S3**). Moreover, sequence variations in dsRNA compactness have been found to be consistent for different force-field and water model choices (32) (33). In addition to the helical rise, other structural parameters such as inclination and roll were found to be highly negatively correlated with the major groove width (**Figs. S4, S5**) in agreement with (33).

### AU-tracts induce a curvature in dsRNA by local compression of the major groove

We have shown that the sequence of homogeneous dsRNA molecules modulates the major groove size, with the poly-AU sequence leading to a significant compaction of the major groove width. Compaction of the minor groove in DNA A-tracts has been linked with the intrinsic bending induced by these sequences (23). Motivated by the DNA case, we next explored if major groove narrowing could lead to bent structures in dsRNA. In order to do so, the poly-G sequence, which showed standard values of the structural parameters (**Fig. 1**), was modified to include a stretch of alternating A’s and U’s, hereafter AU-tract. We thus simulated five different sequences with AU-tracts of lengths varying from three to seven base pairs, which were denoted as AU-3 to AU-7 (**Table 1**). All sequences were designed to be similar in length (24 or 25 bp) and to contain the AU-tract exactly in the center of the duplex.

Our results revealed a localized compression of the major groove at the position of the AU-tract. This can be seen in **Fig. 2a**, where we represented the major groove width profiles of the poly-G (same as **Fig. 1a**), AU-4 and AU-6 sequences. The homogeneous major groove width of the poly-G molecule (**Fig. 2a** left) contrasted with the abrupt drop found when a 4 bp-long AU-tract is introduced (**Fig. 2a**, middle). As the length of the AU-tract increased, the reduction of the major groove width was more pronounced, reaching lower values and extending over longer distances along the duplex. This effect can be appreciated in the major groove profile of AU-6, which presented a deeper and wider drop than that of AU-4, reaching a minimum value lower than 5 Å (**Fig. 2a**, right). Notice that a short 3 bp-long AU-tract was enough to induce major groove narrowing, although the effect was amplified in longer AU-tracts (**Fig. S6**).

**Figure 2.**
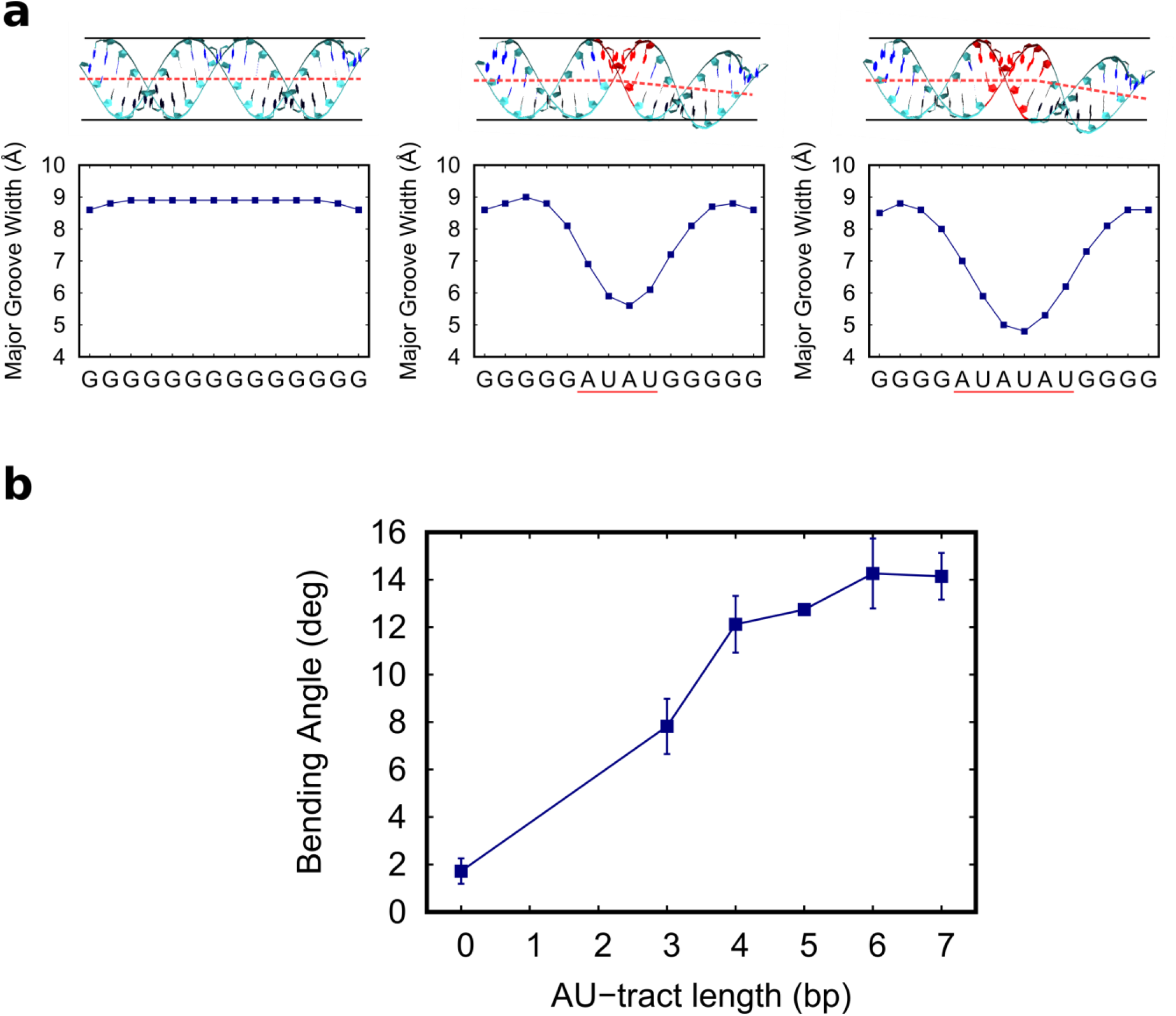
AU-tracts produce a bend when inserted in poly-G dsRNA molecules. **a**, Top: average structure of the poly-G, AU-4 and AU-6 sequences with the AU-tracts highlighted in red. The black lines represent a cylinder, which is unable to embed highly bent molecules, namely AU-4 and AU-6. An approximate helical axis was drawn in red dotted line to guide the eye. **a**, Bottom: major groove width profiles of poly-G, AU-4 and AU-6 were computed and represented as in **Fig. 1a**. Localized drops in these profiles are found in the AU-tracts (underscored in red) which coincide with the bending region of the molecule (top). **b**, Bending was computed for the AU-tracts of different lengths (values of the AU-tract length sequences are shown in **Table 1**). We divided the trajectories into five 200 ns-long windows and we then computed the average structure over each of these sub-trajectories. The bending angle of these average structures was then obtained using the curvilinear helical axis from the software Curves+ (40) and neglecting four base pairs on each terminal of the molecule. The plotted points are the mean values of the five time windows and the errors are the standard deviations. A line connecting the points was drawn to guide the eye.

Interestingly, compression of the major groove by AU-tracts resulted in bent dsRNA structures. This can be noticed by visual inspection of the computed average structures of the molecules throughout the simulation time. The average structures of AU-4 and AU-6 presented a bend at the position of the AU-tract and were therefore unable to be embedded inside a virtual cylinder. The same reasoning applies to the other AU-tracts, namely AU-3, AU-5 and AU-7 (see **Fig. S6**). On the contrary, the poly-G was straight and, therefore, could be fitted inside a cylinder (see **Fig. 2a**). In order to provide a quantitative description of this bending we resorted to the curvilinear axis definition of Curves+ (40). We first divided our μs-long simulations into five windows of 200 ns and computed the average structure of these subtrajectories. For each of these structures, we computed the bending angle, discarding the four base pairs adjacent to each molecule’s termini. We then averaged over these five measurements for each of the sequences. This bending angle was plotted as a function of the AU-tract length in **Fig. 2b**, where we also included the poly-G homopolymer, which can be considered as a zero-length AU-tract. These measurements corroborated the bending effect of AU-tracts that we inferred from visual inspection of the dsRNA structures. The poly-G sequence, which lacks AU-tracts, was found to be essentially straight, as quantified by a very small bending angle of ~2 deg. Interestingly, the shortest AU-tract considered, which was only 3 bp long, already induced a significant bending of ~8 deg in the RNA duplex. This value increased with the AU-tract length, saturating at ~14 deg with AU-tracts of 6 bp or longer.

### AU-tracts are a major source of bending in random dsRNA sequences

The homogeneous sequences – repeating dinucleotides or mononucleotides – studied so far have allowed us to unveil the phenomenon of AU-tract bending and to relate the length of the tract with the magnitude of the bending. We next explored whether AU-tract bending occurs in the context of heterogeneous sequences by performing a new set of simulations of 24 bp-long arbitrary dsRNA sequences containing an AU-tract (**Table 1**, highlighted in red) as well as other potential bending motifs (**Table 1**, highlighted in green). As in previous sections, simulations were extended to 1 μs time and bending was evaluated from the average structures discarding the four terminal base pairs from each side.

Importantly, only those sequences containing AU-tracts, namely Seqs. 1-3, were significantly bent, showing values of the bending angle larger or equal than 10 deg (**Fig. 3**). Among these three sequences, Seq. 1, which has the shortest AU-tract – 3 bp long – scored the lowest bending angle, but still substantially larger than any of the sequences lacking AU-tracts (Seqs. 4-6). Consistently, Seq. 1 presented a less pronounced drop in the major groove width compared with Seq. 2 and Seq. 3, which contain a longer AU-tract comprising four base pairs. Seq. 4 and Seq. 5 contained no AU-tract, but other motifs rich in A’s and U’s, namely AUU and UAA. Contrary to AU-tracts, these motifs produced very modest variations in the major groove width and, consequently, Seq. 4 and Seq. 5 were nearly straight (**Fig. 3**). Seq. 6 presented a CG-tract that locally enlarged the major groove, in line with the results from **Fig. 1a**. However, this effect was not translated into an enhanced bending of the duplex.

**Figure 3.**
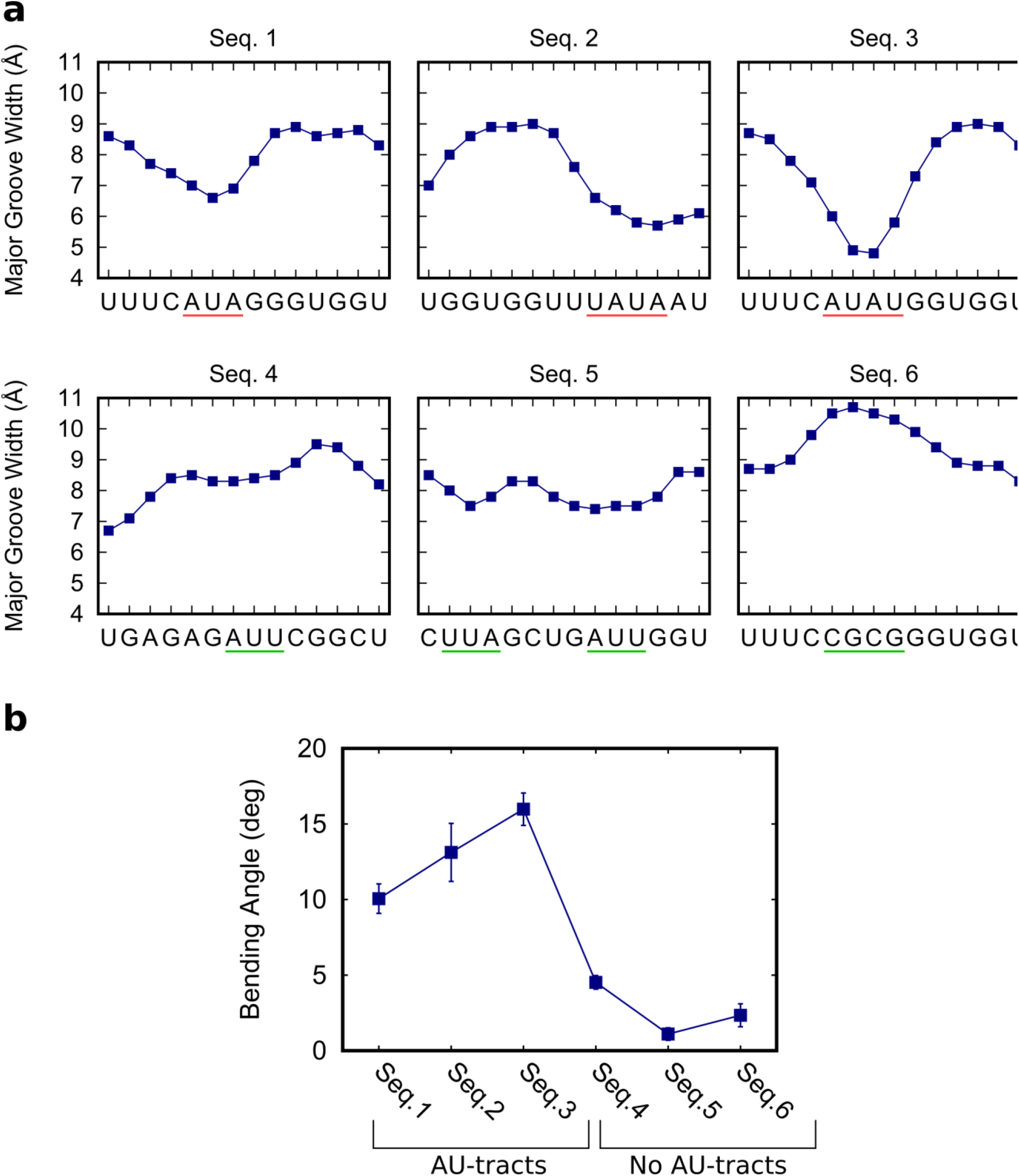
AU-tracts bend arbitrary dsRNA sequences. **a**, Major groove width profiles of the test sequences (see **Table 1**) were computed and represented as done in **Fig. 1a**. AU-tracts are highlighted in red and other tested motifs, such as AUU or CGCG, are highlighted in green. **b**, The bending angle was measured for the arbitrary sequences as done in **Fig. 2b**. Only Seqs. 1-3, which contain an AU-tract, scored a bending angle larger or equal than 10 deg.

These results are in line with crystallographic studies reporting a bend in duplexes containing a central AU-tract (30) (31) (43). However, crystal packing can induce spurious bending in nucleic acids (44) and, indeed, the bent helices observed in the AU-tract structures were partly attributed to intermolecular interactions among different duplexes of the crystal (30) (31). In the following section we experimentally demonstrate that AU-tracts promote the formation of bent dsRNA structures at the single-molecule level.

### Atomic force microscopy imaging shows that AU-tracts bend the RNA duplex

Motivated by our simulation findings, we performed atomic force microscopy (AFM) imaging to experimentally test the effect of AU-tracts on dsRNA bending. We hypothesized that AU-tracts located in phase with the dsRNA helical pitch would amplify their bending, similar to the case of A-tracts in hyperperiodic DNA sequences (18) (21) (24) (45). Therefore, we synthesized two dsRNA constructs that contained phased repetitions of an AU-tract with a periodicity of 11 bp. The first of these constructs was 612 bp-long and contained a periodic AU-tract of 4 bp; the second one, comprised 624 bp and the periodic AU-tract was 5 bp in length. These molecules were correspondingly denoted as ExpAU-4 and ExpAU-5 (see Table 2 and **Supplementary Material**). As control, we considered an arbitrary dsRNA sequence of 612 bp and GC-content of ~ 50% (see **Table S1**). **Figure 4a, b and c** show representative AFM images of control and AU-tracts dsRNA molecules. From the AFM images, we measured the contour length of the molecules and obtained a value of 179 nm for the three constructs, with an error (standard error of the mean) of 3 nm for the control and 4 nm for both AU-tracts molecules (see **Table 2**). These values yielded a ratio of 2.9 Å/bp, which coincides with crystallographic data of dsRNA (41) and with our MD simulations (**Fig. 1b**). Further details on the preparation of dsRNA molecules and AFM imaging conditions can be found in the **Materials and Methods section**.

**Figure 4.**
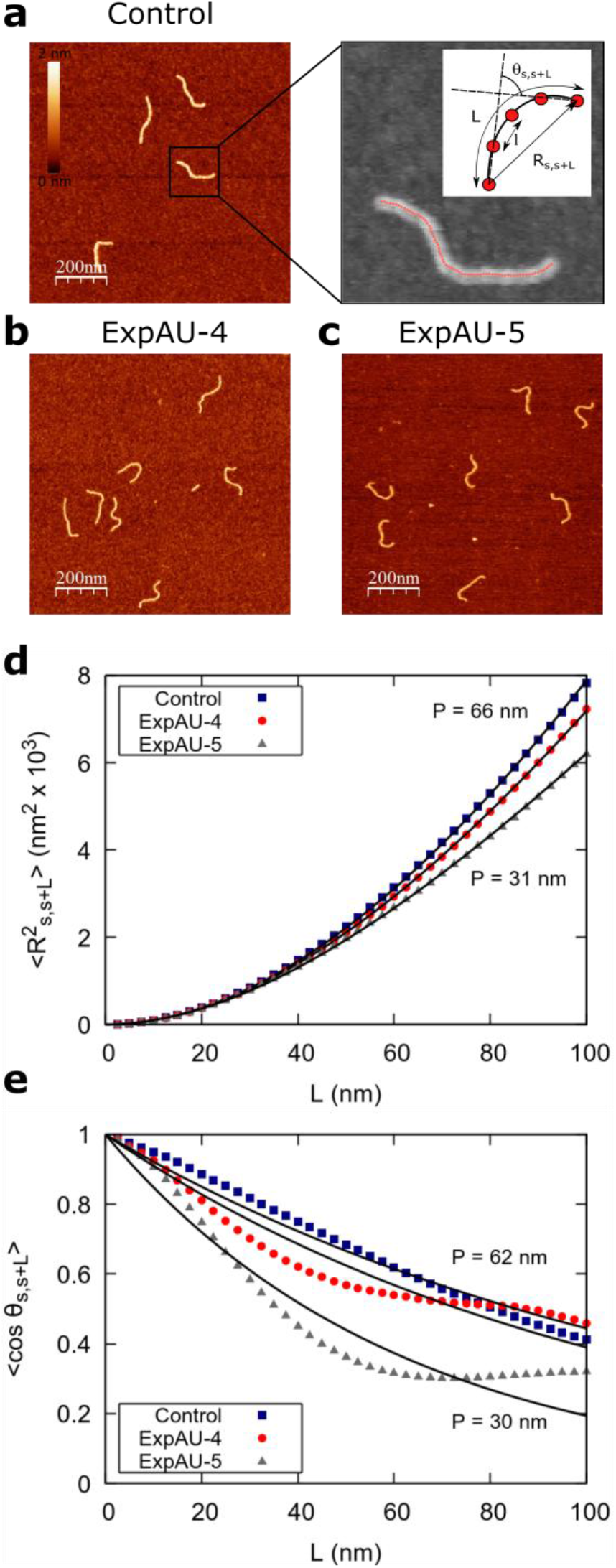
Experimental demonstration of AU-tract bending in dsRNA. **a**, Representative AFM image of control molecules with a zoom-in image of the marked region showing an example of a typical trace. Additional AFM images are shown in Fig. S7. Inset, cartoon depicting a segment of a trace. We represented the end-to-end distance, *R*_*S,S*+*L*_, between two points separated by a contour distance *L* = 10 nm and the angle defined by the tangents at those points, *θ*_*S,S*+*L*_. The distance between two adjacent points of the trace is *l* = 2.5 nm. **b**, Representative AFM images of ExpAU-4 and **c**, ExpAU-5 molecules. Z-scale is the same as in **a**. **d**, Mean squared end-to-end distance plotted as a function of the contour distance between two points. The lines are fits to **Eq. 1**, which yielded *P* =66 ± 1 nm, *P* =46 ± 1 nm and *P* =31 ± 1 nm for the control, ExpAU-4 and ExpAU-5, respectively. **e**. The mean cosine of the tangents was plotted as a function of the contour distance between two points. The control data approximately fitted to the WLC model with *P* =62 ± 1 nm. The AU-tracts data showed significant deviations from the WLC and the best fit yielded a persistence length of *P* = 53 ± 1 nm for the ExpAU-4 and 30 ± 1 nm for the ExpAU-5.

**Table 2.** Experimental AFM measurements on AU-tract dsRNA bending. The control was an arbitrary dsRNA sequence of 612 bp; ExpAU-4 and ExpAU-5 sequences consisted on periodic repetitions of a 4 bp-long and a 5 bp-long AU-tract, respectively. The letter N denotes an arbitrary nucleotide. Sequences are written from the 5’ to the 3’ end, omitting the complementary strand. The full sequences can be found in **Table S1**. The contour length showed a similar value for the three molecules. The persistence length was obtained by fitting the mean squared end-to-end distance (< *R*^2^ >, fourth column) and the cosine’s correlation (< cos *θ* >, fifth column) to the WLC model (**Eq. 1** and **Eq. 2**, respectively). Errors in the contour length are the standard error of the mean and errors in the persistence length are the ones from the fit.

| Label | Sequence | Contour length | $P$ (nm) from $\langle R_{S,S+L}^2 \rangle$ | $P$ (nm) from $\langle \cos \theta_{S,S+L} \rangle$ | Number of molecules |
| --- | --- | --- | --- | --- | --- |
| Control | Arbitrary, 612 bp | 179±3 | 66 ± 1 | 62 ± 1 | 202 |
| ExpAU-4 | N<sub>7</sub> (AUAU N<sub>7</sub>)<sub>55</sub> | 179±4 | 46 ± 1 | 53 ± 1 | 127 |
| ExpAU-5 | N<sub>7</sub> (UAUAU N<sub>6</sub>)<sub>56</sub>N | 179±4 | 31 ± 1 | 30 ± 1 | 195 |

By visual inspection of **Fig. 4** and **Fig. S7** one can already notice that AU-tracts molecules are more bent than the control. In order to quantitatively assess the bendability of the control and AU-tracts dsRNA molecules, we first obtained traces of points separated by *l* = 2.5 nm that follow the trajectory of the molecules (21) (24) (46) (**Fig. 4a**, right). From these traces, we computed the end-to-end distance, *R*_*S,S*+*L*_ between two points separated by a given contour length *L*, and the angle, *θ*_*S,S*+*L*_, defined by the tangents to the trajectory at those points (inset in **Fig. 4a**, right). We then computed the square of *R*_*S,S*+*L*_ and the cosine of *θ*_*S,S*+*L*_ and averaged over all the points of a trace and over all the measured traces. The resulting mean squared end-to-end distance, 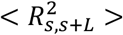 and mean cosine of the tangents, < cos *θ*_*S,S*+*L*_ >, allowed us to study the mechanical properties of dsRNA in the context of polymer physics models, concretely the widely used worm-like chain (WLC) model:

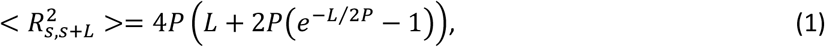

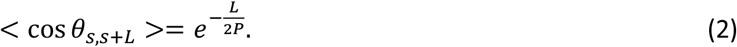

In these equations, *P* is the persistence length, which is directly proportional to the bending rigidity (*B*) of the polymer *P* = *B*/*K*_*B*_*T* (where *K*_*B*_ is the Boltzman constant and *T* is the temperature). When fitted to **Eq. 1**, the data of the control dsRNA yielded a persistence length of *P* = 66 ± 1 nm, consistent with previous single-molecule experiments on arbitrary dsRNA sequences (38) (39) (47) (**Fig. 4d**), and proving that adsorption conditions preserve equilibrium conformations of the polymer. However, the ExpAU-4 molecule presented a persistence length of *P* = 46 ± 1 nm, around 30% lower than that of the control. The ExpAU-5 molecule showed an even lower value of the persistence length, *P* = 31 ± 1 nm, which was around 50% the value of the control. The low values of dsRNA persistence length found for the AU-tract sequences indicated that these molecules were more bendable than arbitrary dsRNA sequences. Moreover, as the length of the periodic AU-tract was increased from 4 bp to 5 bp, the magnitude of the bending also increased, as manifested by the reduction in the persistence length of the ExpAU-5 with respect to the ExpAU-4.

Further analysis of our data in terms of the cosines of the tangents provided a more stringent test to the WLC model (24) (39). For the control molecule, the cosines of the tangents reasonably fitted to the WLC model (**Eq. 2**); for the AU-tracts molecules, on the contrary, they did not (**Fig. 4e**). A fit of the control data to the WLC resulted in a persistence length of *P* = 62 ± 1 nm, slightly lower, but consistent with the value obtained using 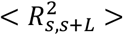 (**Fig. 4d**). The ExpAU-4 molecule presented clear deviations from the WLC model and the best fit yielded a value of *P* = 53 ± 1 nm. The deviations from the WLC behavior were even more significant for the ExpAU-5 molecule, which showed a local minimum of < cos *θ*_*S,S*+*L*_ >= 0.3 at around *L*~ 60 nm. We obtained a value of *P* = 30 ± 1 nm when we fitted the < cos *θ*_*S,S*+*L*_ > data of the ExpAU-5 to **Eq. 2**. The fact that the WLC could not capture the flexibility of AU-tracts molecules is indicative of the existence of an intrinsic bending that is not contemplated by purely entropic models.

### AU-tracts: similarities and differences with DNA A-tracts

Sequence-dependent bending is known to take place in dsDNA by means of A-tracts: sequences of at least four A·T base pairs without a TA step. When several A-tracts are located in phase with the helical pitch they produce a macroscopic curvature in the DNA (23). This curvature can be directly observed using AFM or electron microscopy (21) (24) (48) (49), or can be inferred from gel electrophoresis experiments (18) (50). In addition, A-tracts display a particular conformation at the molecular level, which differs from that of canonical B-DNA (23) (25). In the following, we compare these well-known features of dsDNA A-tracts – macroscopic curvature and molecular conformation – with our findings on dsRNA AU-tracts.

Previous AFM works have provided a detailed picture of bending deformations in dsDNA molecules with A-tract-induced curvature. These experiments showed that, as a consequence of that curvature, the structural properties of dsDNA sequences with phased A-tracts exhibit significant deviations from the WLC model and that the best fit to that model yields a low value of the apparent persistence length (21). These two effects – deviations from the WLC and low *P* – were also observed for the dsRNA AU-tracts (see **Fig. 4d, e**). Although our AU-tracts 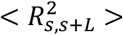 data showed no clear discrepancy with respect to the WLC prediction, such deviations only appeared in the A-tracts for contour lengths greater than ~120 nm length (21) (24), which are beyond the range studied here (<100 nm). The < cos *θ*_*S,S*+*L*_ > of the A-tracts, on the contrary, deviated from the WLC behavior at shorter contour lengths (~50 nm) and is, therefore, a better indicator of the existence of intrinsic curvature. Consistent with the presence of intrinsic bending, our AU-tracts also presented significant deviations from the WLC in the < cos *θ*_*S,S*+*L*_ > (*L*) data. Moreover, the shape of the < cos*θ* > (*L*) plot for the phased AU-tract studied here is remarkably similar to an intrinsically-bent A-tract dsDNA that we recently reported (24). Finally, we checked whether the AU-tracts caused anomalous migration of dsRNA in agarose gels, as occurs for A-tracts in dsDNA molecules. Under the conditions studied, we found that AU-tracts molecules showed no anomalous migration (see **Fig. S9**).

We then turned our attention to the molecular structure of A-tracts and AU-tracts. We thus compared the structural features of the dsRNA AU-tract from Seq. 3 (see **Table I**) with the DNA A-tract from a high-quality NMR structure of the Drew-Dickerson Dodecamer (DDD) (51). The DDD is the most extensively characterized DNA duplex and contains a central A-tract: CGCGAATTCGCG. The comparison of the most relevant structural parameters revealed intriguing similarities and differences between A-tract and AU-tract bending (**Fig. 5**). As expected, the central region of the DDD shows the standard features of A-tracts, which are a highly negative propeller twist, a narrowing of minor groove and a negative roll (52) (53) (54). Moreover, the major groove width showed little variation and all the plots were symmetric, a consequence of the palindromic sequence of the DDD. Similar to the A-tract case, a large negative propeller twist was observed in the AU-tract. However, the roll parameter presented very different trends in the two molecules: a maximum at the center of the AU-tract, but a minimum in the A-tract. This difference in roll can be associated to the changes observed in the dimensions of the grooves. Positive roll values are attributed to bending towards the major groove and, therefore the increase in roll is consistent with the compression of the major groove observed in the AU-tracts (**Fig. 2a** and **Fig. 3a**). Conversely, the high negative roll found in A-tracts can be associated with their narrow minor groove, a well-known feature of these sequences (52) (53) (54). Furthermore, notice that only one of the grooves showed a significant sequence variation. Namely, the minor groove of the AU-tract and the major groove of the A-tract were approximately homogeneous. Consistently, similar trends of these structural parameters were obtained with the alternative analysis software 3DNA (**Fig. S8**).

**Figure 5.**
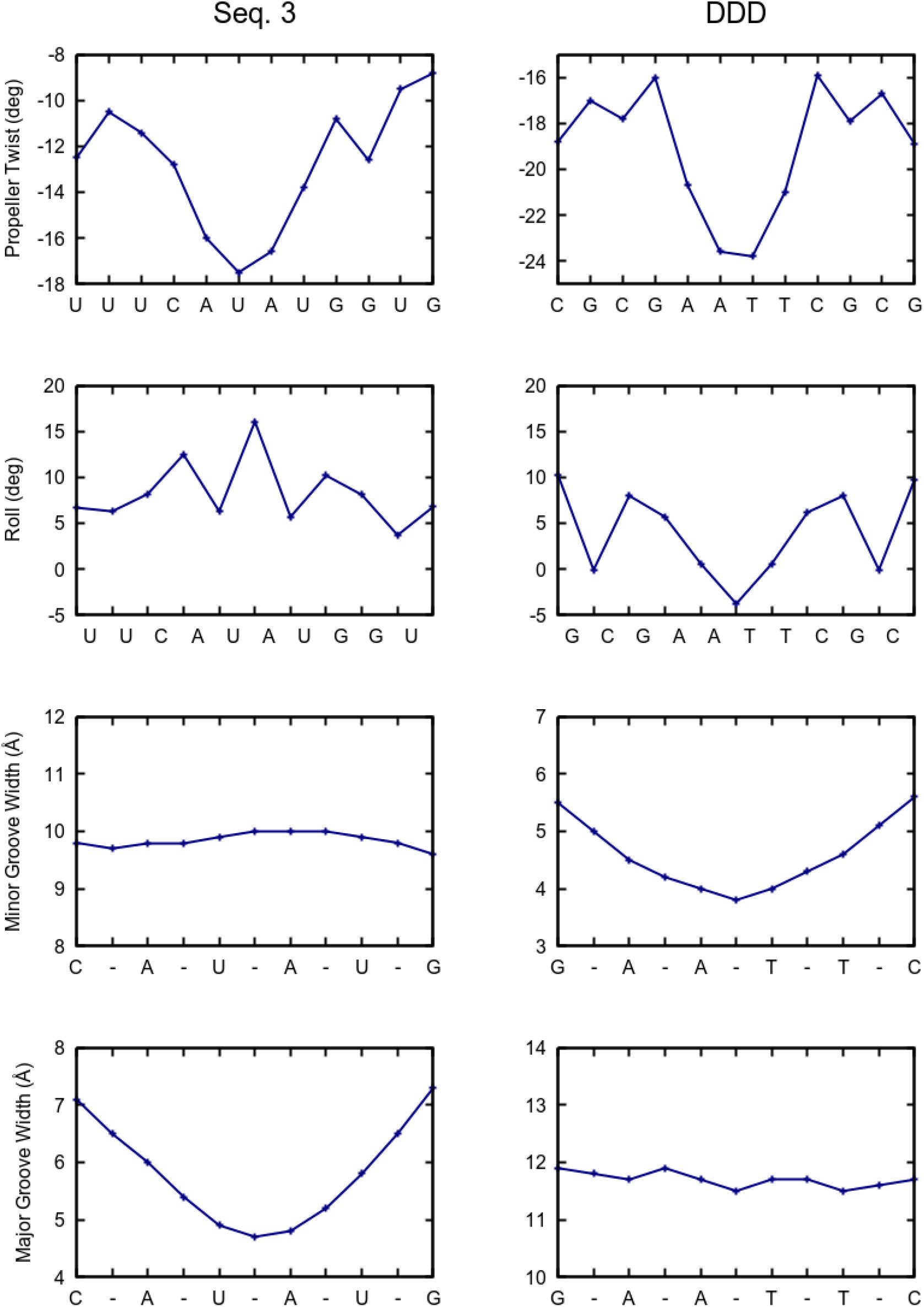
Comparison of structural parameters of dsRNA AU-tracts and dsDNA A-tracts. The average structure of the simulated Seq. 3 and a recent NMR (51) structure of the Drew Dickerson Dodecamer (DDD) were taken, respectively, as representative examples of a dsRNA AU-tract and a dsDNA A-tract structure. The analysis was performed using Curves+ (40). In addition to the major groove width, the structural parameters propeller twist, roll and minor groove width, which are typically used in the characterization of A-tracts (23) were computed.

### Implications for dsRNA recognition

dsRNA structures are ubiquitous in cells, and together with dsRNA-binding proteins are central players in cellular processes, such as mRNA biogenesis and editing, microRNA processing and function, as well as anti-viral defense (55) (3) (56) (57). In some of these processes, dsRNA-binding proteins can be rather selective in their target RNA sequences, although the recognition mechanism is not completely understood (58). Interestingly, these protein-dsRNA complexes sometimes present bent dsRNA structures. For example, dsRNA bending was observed in the crystal structure of an RNA duplex in complex with MDA5 (13) or OAS1 (15). Moreover, bending was predicted to occur in dsRNA upon the interaction with RIG-I (59) or Dicer (14) (60). One can thus speculate that the dsRNA curvature at the AU-tracts observed in the present study might play a role in specific target recognition by dsRNA-binding proteins. Namely, it is conceivable that AU-tracts sequences will be preferred in protein-dsRNA interactions that require dsRNA bending. Finally, the narrow major groove characteristic of AU-tracts might also contribute to achieve dsRNA sequence specificity, as occurs with the minor groove in dsDNA A-tracts (28) (29). This would add to other mechanisms of dsRNA sequence recognition, such as the recently proposed contacts through the minor groove (58) (61).

## CONCLUSION

In this work we have investigated the possible effect of the nucleotide sequence on dsRNA bending. Our molecular dynamics simulations of repeating dinucleotide sequences revealed that the poly-AU sequence adopted a characteristic conformation with a narrow major groove. By inserting AU-tracts of different lengths inside a poly-G homopolymer, we found that a 3 bp-long AU-tract was enough to induce a bend, but longer AU-tracts resulted in larger bending. This finding was consistent when studying AU-tracts located in arbitrary sequences. Finally, our simulation results guided the design of dsRNA constructs suitable for measuring the effect of AU-tract bending in AFM experiments. Using AFM imaging, we found that these AU-tract molecules were more bent than control dsRNA’s of arbitrary sequence, confirming the prediction of our simulations. More importantly, the AU-tracts showed deviations from the WLC behavior, a hallmark of intrinsic curvature.

Intrinsic bending induced by dsDNA A-tracts has been linked to multiple biological functions such as nucleosome positioning, localization of supercoils or germ-line gene silencing. It is therefore expected that the sequence-dependent bending reported here for dsRNA might also have important biological implications. On one hand, the bent structure of the AU-tracts could be exploited in the formation of tertiary contacts in the process of RNA folding. On the other hand, AU-tracts might provide a mechanism for sequence recognition based on dsRNA shape. Hence, our results might shed new light on how dsRNA sequence dictate specificity on dsRNA-protein interactions, and thereby have impact on biological processes ranging from antiviral response to gene silencing.

## Materials and Methods

### Molecular Dynamics Simulations

Simulation details are similar to the ones from (62), only excluding the external force. RNA duplexes were placed in an approximately cubic box of 110 Å edge size and filled with water and sodium counterions to balance the phosphate charges. The systems were heated up to 300 K and equilibrated in the isobaric-isothermal (NPT) ensemble (P = 1 atm, T = 300 K) for 20 ns. Production simulations were run in the NVT ensemble using as input coordinates the ones from the last configuration of the NPT equilibration. All simulations were extended to ~1 μs time.

We used the AMBER14 software suite (63) with NVIDIA GPU acceleration (64) (65) (66). For the modeling of dsRNA molecules we resorted to the Cornell ff99 force field (67) with the parmbsc0 (68) refinement and the χOL3 modification (69). The ions were described according to the Joung/Cheatham (70) parametrization; and the TIP3P model (71) was used for water molecules. Periodic boundary conditions and Particle Mesh Ewald (with standard defaults and a real-space cutoff of 9Å) were used to account for long-range electrostatics interactions. The same real space cutoff was used to truncate Van der Waals forces. SHAKE algorithm was used to constrain bonds containing hydrogen atoms, thus allowing us to use an integration step of 2 fs. Coordinates were saved every 1000 steps. Average structures were computed using the *cpptraj* software of the AMBER14 suite (63). Helical, base pair step parameters and groove dimensions were computed using Curves+ (40) and 3DNA (42) and helical bending was calculated using Curves+ (40). The four base pairs adjacent to the termini of the molecules were excluded from the analysis.

### Production of dsRNA molecules

In order to study the mechanical properties of AU-tracts at the single-molecule level, we produced dsRNA molecules by hybridizing two complementary ssRNAs. For that, the sequence of interest was cloned after the T7 RNA polymerase promoter between two KpnI sites. In this way, the fragment could be digested and ligated in the opposite orientation, allowing us to synthesize the two complementary ssRNA chains. In addition, a SmaI site was introduced at the end of the sequence, enabling the linearization of the plasmid vector that was used as transcription template.

We first produced the ExpAU-4 dsRNA molecule (**Table 2**). The first step was to produce a plasmid by ligation of two gel extracted (QIAGEN) DNA fragments: one from the pBlueScriptIISK+ plasmid (Stratagene) and the other from the pNLrep plasmid. This plasmid was used to obtain a set of plasmids by several rounds of ligation of different pairs of hybridized oligonucleotides, in such a way that the resulting DNA segment always contained repetitions of the ATAT sequence separated by seven random base pairs (ATAT)N_7_. In the first ligation, we employed the oligonucleotides *98.5P-F-T7* and *99.5P-R-T7* (fragment 98 and 99, **Table S2**) in order to locate the T7 promoter before the AT-periodic region. We then performed five additional rounds of ligations in different restriction sites of the hybridized oligonucleotides *100.5P-F-XhoI-blunt* and *101.5P-R-XhoI-blunt* (fragment 100 and 101, **Table S2**), *102.5P-F-blunt-blunt* and *103.5P-R-blunt-blunt* (fragment 102 and 103, **Table S2**). After six rounds of ligation, the plasmid (pBlueSK-T7-6oligos) contained a DNA segment with the final length. A similar strategy was followed to produce the ExpAU-5 dsRNA molecule. In this case, we fabricated a set of plasmids that contained a segment of repetitions of the TATAT sequence separated by six random base pairs TATAT(N)_6_. The three pairs of hybridized oligonucleotides employed here were: *121.5P-F-T7* and *122.5P-R-T7* (fragment 121 and 122, **Table S2**), *123.5P-F-XhoI-blunt* and *124.5P-R-XhoI-blunt* (fragment 123 and 124, **Table S2**), *125.5P-F-blunt-blunt* and *126.5P-R-blunt-blunt* (fragment 125 and 126, **Table S2**). Finally, a control dsRNA with a GC content of ~50% and the same length of the ExpAU-4 molecule (612 bp) was produced. To fabricate this molecule, we PCR amplified a Lambda DNA fragment (NEB) of ~50% GC-content with the oligonucleotides *114.F RNA control 612* and *113.R RNA control 1316*. The PCR product was digested, purified and ligated with the long dephosphorylated fragment purified after digestion of pBlueSK-T7-6oligos plasmid with KpnI. All the plasmids were checked by DNA sequencing.

dsRNA molecules were synthesized according to a previously described protocol (39) (72) with slight modifications to increase yield for single-molecule manipulation purposes. Once each pair of plasmids were obtained, plasmid vectors used as transcription templates were linearized with SmaI followed by purification (QIAGEN). Afterwards, *in-vitro* transcription using the commercial HiScribe™ T7 High Yield RNA Synthesis Kit (NEB) gave rise to two complementary ssRNAs without any non-complementary nucleotides at their ends. After 3 h at 42 °C, EDTA was added to a final concentration of 30 mM, and both strands were subsequently hybridized by heating 1 h at 65 °C and slowly cooling to room temperature at a 1.2 °C/5 min rate up to 25 °C. This resulted in dsRNA molecules without any single-stranded overhangs. Transcription products were then cleaned with RNeasy MinElute Cleanup Kit (QIAGEN) followed by treatment for 1 h at 37 °C with 2.5 units of RNase-free DNase I (NEB). The sample was once again cleaned with RNeasy MinElute Cleanup Kit before applying on a 1% agarose gel for gel extraction and purification with QIAGEN gel extraction kit and elution with RNase free H_2_O. dsRNA constructs were stored at 4 °C in RNase free H_2_O. The final sequences are shown in **Table S1**.

### Atomic Force Microscopy Measurements

Imaging conditions and data analysis are similar to those employed in a previous work (39). A 10 μL solution containing 0.5 nM dsRNA, 2.5 mM NiCl_2_, 25 mM TrisAc pH 7.5, 2.5 mM MgOAc and 100 mM NaCl was deposited onto freshly cleaved mica. After ~60 s, the sample was washed using Milli-Q water and dried using air nitrogen. Images were taken in tapping mode in air, using an AFM from Nanotech Electronica S.L. with PointProbePlus tips (PPP-NCH Nanosensors). Contour lengths were obtained using the WSxM software (73). Persistence lengths were computed using the tracing routine from (21) (46). Traces of 170 nm were obtained with a point-to-point separation of 2.5 nm.

## ACKNOWLEDGEMENTS

We thank the financial support from the Spanish MINECO (projects MAT2017-83273-R (AEI/FEDER, UE) to R.P., BFU2017-83794-P (AEI/FEDER, UE to F. M.-H.) and Comunidad de Madrid (Tec4Bio – S2018/NMT-4443 and NanoBioCancer – Y2018/BIO-4747 to F.M.-H.). R.P. acknowledges support from the Spanish Ministry of Science and Innovation, through the “María de Maeztu” Programme for Units of Excellence in R&D (CEX2018-000805-M). F.M.-H. acknowledges support from European Research Council (ERC) under the European Union Horizon 2020 research and innovation (grant agreement No 681299). J.G.V. acknowledges funding from a Marie Sklodowska Curie Fellowship within the Horizons 2020 framework. Alberto M.-G. acknowledges support from the International PhD Program of “La Caixa-Severo Ochoa” as a recipient of a PhD fellowship. The authors acknowledge the computer resources, technical expertise and assistance provided by the Red Española de Supercomputacion at the Minotauro Supercomputer (BSC, Barcelona). U.F.G. acknowledges support from the Swiss National Science Foundation (31003A_179256 / 1). Alejandro M.-G. acknowledges support from the Spanish Ministry of Competitiveness and Industry as a recipient of a FPI fellowship (REF BES-2015-071244).

## Contributions of authors

U.F.G. motivated Alberto M.-G., J.G.V., F.M.-H. and R.P to study the sequence-dependent properties of dsRNA. Alberto M.-G., J.G.V., U.F.G., F.M.-H., and R.P. designed research. Alberto M.-G. and J.G.V. performed and analyzed MD simulations. C.A.-R. designed and fabricated the dsRNA molecules for the AFM experiments. Alberto M.-G., M.M.-B. and Alejandro M.-G. performed and analyzed the AFM experiments. U.F.G., M.S., and A.K. suggested test dsRNA sequences based on their experimental work. All authors contributed to discussion of data and review and editing of the manuscript. Alberto M.-G. wrote the original draft of the paper.

